# A Spatio-temporal Investigation of Dynamics of a Two-dimensional Multi-scale Fitz-Hugh Nagumo Neuronal Network

**DOI:** 10.1101/2023.11.10.566630

**Authors:** Alireza Gharahi, Majid Mohajerani

## Abstract

The multi scale architecture by Breakspear and Stam [2] introduces a framework to consider the dynamical processes specific to a nested hierarchy of spatial scales, from neuronal masses to cortical columns and functional brain regions. They hypothesize that the neural dynamics is a function of the structural properties of the neural system at a certain scale as well as the emergent behaviour of the smaller scale activities. In this paper, we adopt the multi scale framework to investigate a generalized version of the stochastic Fitz-Hugh Nagumo (FHN) neuronal system within the small scale process and their emergent large scale synchronization effects leading to the formation of travelling waves in the large scale system. We extend the multi scale framework to incorporate the nonlinear biological synaptic connectivity at the neuronal mass scale. The modified multi scale scheme utilizes the two-dimensional wavelet decomposition in the plane of dynamical interconnected neurons. In addition, we consider the large-scale spatio-temporal system of FHN reaction-diffusion partial differential equations and evaluate the formation of travelling waves in the simplified context of a cellular neural network (CNN) model. Numerical examples are given to illustrate the response and the isolated influence of the strength of neural connectivity on the travelling wave formation modes.

## 1 Introduction

Theoretical and computational models of neural activities aim to enhance the foundation of our understanding of the nature of the neural systems, the analysis and interpretation of the neuroscience functional data. These models, in essence, elaborate the patterns of spatio-temporal neural activity by replicating the idealized underlying dynamical mechanisms. Further, neuronal models instruct the design of empirical techniques and regulate the corresponding algorithms. To this end, computational approaches that best suite the research into cognition, perception and motor activities are directed toward the collective behaviour of many neuronal populations rather than single spikes. However, establishing a theoretical framework that gives rise to enhanced mathematical models of large scale collective activities remains a challenge due to the variety of technical and conceptual limitations [3].

A reasonable approach to the neural dynamics problem has to incorporate individual highly nonlinear spikes of neurons, the synaptic interaction of the individual neurons, and the random processes involved in the dynamical state of the system which altogether lead to a high-dimensional representation. Such a complex system requires a scheme that can reduce the intractable dimensions to bridge between the small scale patches of cortical populations dynamics and the large-scale neural activities of the type that are observed in macroscopic imaging data such as functional magnetic resonance imaging (fMRI) and electroencephalography (EEG) [4, 17]. This is achieved by coupling dynamics of neural mass ensemble circuits into a hierarchy of scales each with their specific neural connectivity characteristics. In this respect, Bearkspear and Stam [2] proposed a wavelet-based multi-scale setting which incorporates a set of evolution equations of physiological processes into the coupling structure with a formalism that allows the modular architecture of repeated neuropil over a hierarchy of scales, simulating the brain organization. In their work, the authors present a systematic construct that incorporates the local dynamics through the neurobiological processes specific to each scale along with the coupling rule (connectivity of neural ensemble) corresponding to that scale and a multi scale hierarchy of such ensembles with between scale reciprocalities. In the proposed construct they utilize the orthogonal wavelet decomposition to project spatio-temporal information from one scale to another by the appropriate spatial variance, and at the same time, preserve the localized character of the dynamics in the between scale projection. Breakspear and Stam [2] apply the provided multi scale framework to a Morris-Lecar system [15] and demonstrated the inter-scale dependence based on the variety of model properties; a model realization of a one-dimensional signal arrays with linear within-scale and inter-scale couplings. In the present work, we would like to examine the Breakspear-Stam [2] scheme in the context of the well-known FitzHugh-Nagumo (FHN) model within biologically determined stochastic nonlinear synaptic couplings at the small scale view, and the reaction diffusion type equations across the neural domain at the large-scale. We investigate three categories based on the connectivity formation of the underlying neural network at the small scale; fully connected, moderately connected, and weakly connected network. Our additional contribution is to extend the multi scale layout to signal decomposition on a spatially two dimensional neural network and to demonstrate the emergence of travelling waves in the large scale system as a result of the connectivity strength of the small scale stochastic neuronal dynamics. Finally, we examine the criteria for the presence of large scale periodic solutions and emergence of travelling waves within a simplified deterministic variant of the model.

## 2 Preliminaries

### 2.1 Wavelet decomposition transform

In this preliminary section, we provide a brief description of the concept of the wavelet multi-resolution method in multi-scale analysis [24]. The multi-resolution approach to multi-scale analysis considers a sequence of nested subspaces of the ambient space, e.g the space of square-integrable functions, *L*^2^(ℝ). Let us denote these subspaces by {*V*_*j*_, *j* ∈ ℤ} each containing functions with components up to the *j*th scale. The sequence of subspaces {*V*_*j*_}_*j*∈ℤ_ is a multi-resolution if, for all *j* ∈ ℤ *V*_*j*_ ⊂ *V*_*j*+1_; the subspaces are self-similar in scale, i.e. a function *f* (*x*) ∈ *V*_0_ if and only if *f* (2^*j*^*x*) ∈ *V*_*j*_ (in other words, *V*_*j*_ is a scaled version of *V*_0_); the subspaces are nested, i.e. {0} ⊂ …⊂ *V*_−1_ ⊂ *V*_0_ ⊂ *V*_1_ ⊂ *V*_2_ … ⊂ *L*^2^(ℝ), ∩_*j*∈ℤ_*V*_*j*_ = {0} and 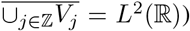 finally the model subspace *V*_0_ is spanned by an orthonormal basis generated by an integer shift of a function *ϕ* known as the scaling function. The orthonormal basis then can be formalized as {*ϕ*(·− *n*), *n* ∈ ℤ}. In most cases *ϕ* requires to be piecewise continuous with compact support. With these four characteristics we define an orthonormal basis for *V*_*j*_ as

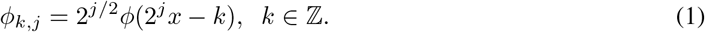

The orthonormal property of the basis {*ϕ*((·− *n*), *n* ∈ ℤ} of *V*_0_ and the fact that *V*_0_ ⊂ *V*_1_ imply the following relation with the orthonormal basis through a sequence of refinement mask coefficients {*a*_*k*_}_*k*∈ℤ_,

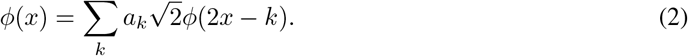

Let *W*_0_ be the orthogonal complement of *V*_0_ in *V*_1_, i.e. *V*_1_ = *V*_0_ ⊕ *W*_0_ and *V*_0_ ⊥ *W*_0_. We define a new function called the wavelet function

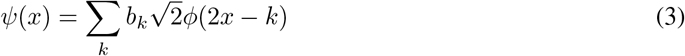

where *b*_*k*_ = (−1)^*k*^*a*_−*k*+1_ for *k* ∈ ℤ. The sequence {*b*_*k*_}_*k*∈ℤ_ is called the wavelet mask. From the orthonormal basis of *V*_0_, {*ϕ*((·− *n*), *n* ∈ ℤ}, it follows that {*ψ*((·− *n*), *n* ∈ ℤ} forms an orthonormal basis for *W*_0_. We similarly introduce the complement subspaces *W*_*j*_ with the aforementioned multiresolution properties, such that

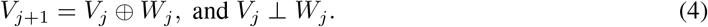

By iterating this decomposition, we can write

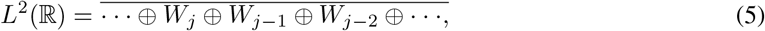

Therefore, *L*^2^(ℝ) can be orthogonally decomposed into {*W*_*j*_}_*j*∈Z_ with the orthonormal basis,

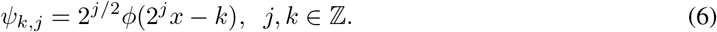

Consequently, for a fixed *j* the sequence {*ψ*_*k,j*_(*x*)}_*j,k*∈ℤ_ forms an orthonormal basis for *W*_*j*_ and the whole {*ψ*_*k,j*_(*x*)}_*j,k*∈ℤ_, an orthonormal basis for *L*^2^(ℝ). Thus, the decomposition principle is founded on construct-ing a wavelet basis generated by the refinable function as discussed in this section [24].

### 2.2 FitzHugh-Nagumo Model

The well known Hudgin-Huxley model [11] (H-H model) is an important classical conductance-based model used computational neuroscience that determines the activity of a single neuron based on the persistent potassium ion K^+^ current through structural changes of four subunit channels, the fast Na^+^ current with three activating gates and one inactivating gate, and a leaking current. The Hodgkin and Huxley established the equation of conservation of electric charge with somatic memberane charge *v*, as follows,

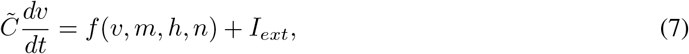

where *m, n*, and *h* are the probability of activating K^+^ gates, activating Na^+^, and inactivating Na^+^ gate being open. In equation (7), 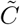 indicates the capacitance and *I*_*ext*_ describes any stimulated current and,

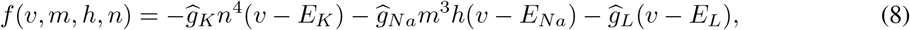

where 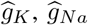, are the maximum conductances of potassium and sodium channels, respectively, and 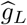 is the conductance of the Ohmic channel. Here, *E*_*K*_, *E*_*Na*_, and *E*_*L*_ are the corresponding Nerst potentials (reversal potentials). The probability of activities can be computed by the following set of linear kinetic ordinary differential equations,

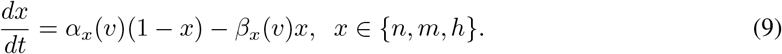

Here, 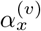 and 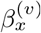 are smooth functions of the membrane potential *v*, and are determined by fitting to experimental data. Obviously, the H-H model consists of a set of four differential equations with four functions to be determined. Many simplifications have been proposed to reduce the H-H model to involve a fewer unknown functions. In particular, FitzHugh and Nagumo reduced the H-H model to an approximating system with two unknown functions for a single neuron [8][7][9][16]. FitzHugh and Nagumo developed a two-dimensional (in the sense of having two unknown functions) model (FHN) of an excitable membrane from Van der Pol’s cubic equation by the addition of terms to guarantee monostability of the system [9]. The model consists of two equations governing the membrane voltage *v* (fast variable) and a recovery variable *w* (slow variable). The voltage equation has a cubic nonlinearity while the slow recovery variable is governed by a linear equation. Involving spatial dependence in the FHN model equations gives rise to a nerve impulse traveling wave solution for the resulting reaction-diffusion system of equations. The spatio-temporal FHN equations take the form,

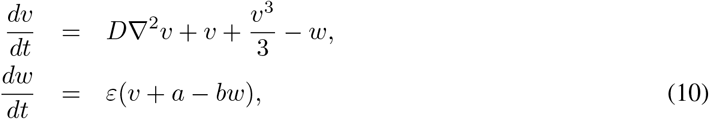

where *v* and *w* are functions of *x* ∈ ℝ and *x* ∈ ℝ^+^, respectively, ∇^2^ indicates the spatial Laplacian operator, and the constants *b*, and *ε* satisfy the conditions: *b* ≥ 0, and 0 *< ε* ≪ 1. Equation (10a) is a nonlinear reaction-diffusion partial differential equation. To construct a network of FHN neurons resembling a neural population we consider a two dimensional spatial grid of *N* ×*N* connected neuronal unites, with connectivity states to be defined.

## 3 Multi-scale Formalism of Neuronal Network

We adopt the wavelet based multiscale architecture of Breakspear and Stam [2] in which the dynamical variables ***u***^(*j*)^(***x***, *t*) are defined across a hierarchy of spatio-temporal scales *j* = *j*_1_, *j*_2_, …, *j*_*l*_. Specific to each hierarchy of scales on evolution equation governs the behavior of the dynamical variable ***u***^(*j*)^(***x***, *t*) and the between scales interdependencies are introduced through the wavelet decomposition. For the purpose of self containment, we revisit the principles of the wavelet transform scheme proposed in the setting of spaciotemporal dynamical systems. In the spatio-temporal modeling of neuronal dynamics, we are intersted in the time evolution of *M* physiological dynamic variables across a spatial domain ***x*** ∈ ℝ^2^ that comprises the components of a vector variable 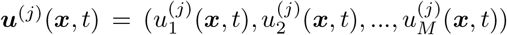. Each component can specify a variable such as membrane potential, cell firing rate, ion channel conductance, etc. Moreover, we modify the setting to accommodate the desired two-dimensional spatial characteristics. As regarded by Breakspear and Stam [2], at each instant *t*, the output of the system is a spatially varying signal ***u***^(*j*)^(***x***, *t*) whose “multi-resolution decomposition” constitute an orthogonal representation of ***u***^(*j*)^ across a specific scale *j*. In principle, we employ the multi-resolution property of the orthogonal wavelet system. Consequently, we construct the two-dimensional wavelet system using the set of scaling functions 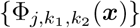 containing information about the fluctuation in the signal of interest at the scale of interest and the wavelet functions 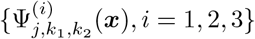 defined by

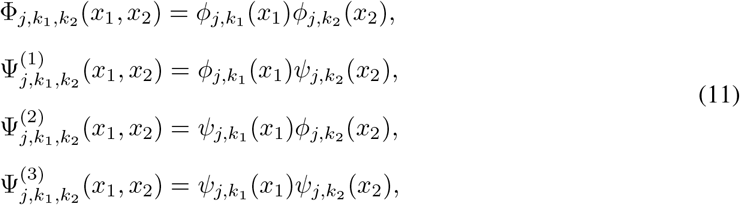

where the spacial coordinates is identified in a Cartesian frame as ***x*** = (*x*_1_, *x*_2_). The multi-dimensional signal in the two-dimensional space ***u***(***x***) can be decomposed at a certain scale *J*, into the sum of the details at the desired scale and the smaller scales and the residual signal at that scale. Accordingly, for any component of ***u***(***x***), i.e. *u*(***x***),

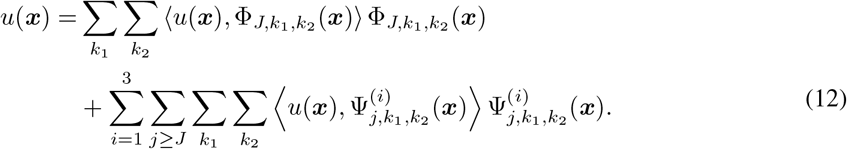

where the first term is the residual approximation and the second term is the superposition of details of scales smaller or equal to *J*. Here ⟨ ., .⟩ denotes the inner product in the *L*^2^(ℝ^2^) space defined by

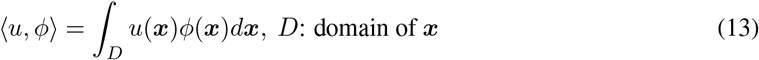

In the current two-dimensional setting, the wavelet functions 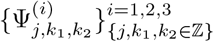 at the scale *j* span the detail subspace *W*_*j*_ as follows,

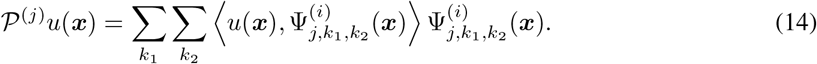

The spatially varying signal *u*(***x***), therefore can be decomposed into an anorthonormal representation across a hierarchy of spatial scales *j* = 0, 1, 2, …, where at each scale *J*, the signal *u*(***x***) is projected onto detail subspaces *W*_*j*_ (*j* ≥ *J*) and an approximation subspace *V*_*J*_ as shown in eqn (12). As in Breakspear and Stam [2005], here we construct the multiscale hierarchy for the spatial dimensions while we establish the temporal dynamics by assembling the spatial projections at each time point. In this paper, we form the multi-scale construct by employing the set of stochastic Fitz-Hugh-Nagumo (FHN) equations for a smaller scale representation of interconnecting neurons.

### 3.1 Small-scale dynamics of interacting neurons

In fact, in the present model the small scale dynamics of single neuronal units are coupled with each other through a nonlinear chemical synaptic interaction model formulated in Baladron et al [1]. The set of stochastic FHN ordinary differential equations governing each neuronal unit is described as

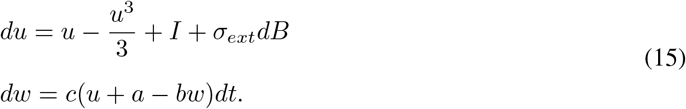

In equation (15), *B* is the standard Brownian motion; *I* is the input current received by the neuron; *a, b* and *c* are recovery constants. A set of coupled equation of the form (15) is established to represent the dynamics of a network the formulated neuron. The inter-neuronal connection in the network is modeled based on the physiological process of the neurotransmitters release from the presynaptic neuron in the synaptic cleft and the subsequent binding to the post-synaptic receptors, what is called chemical synaptic rule whose dynamics is approximated by a first order kinetic scheme. Accordingly, the synaptic current induced by the neural synapses from the presynaptic neuron *i* to the post-synaptic neuron *j* is described as [1]

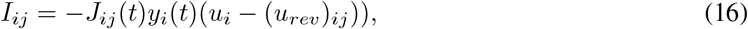

where the synaptic reversal potential (*u*_*rev*_)_*ij*_ is considered a constant, *J*_*ij*_ denotes the maximum conductance and *y*_*i*_ denotes the fraction of open channels commonly considered to be ruled by the following stochastic ODE:

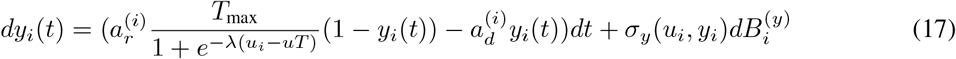

In eq. (17), 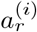 and 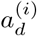 respectively denote the rise and decay rates of the synaptic conductance of the *i*th neuron which for a neural population are commonly considered as constants (*a*_*r*_ and *a*_*d*_). The Brownian motion 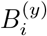 is independent for each neuron *i* and,

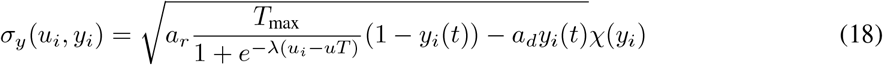

where,

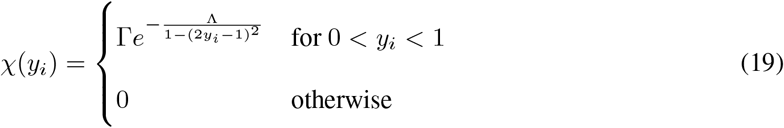

with Γ = 0.1 and Λ = 0.5 taken for all the ionic channels as in [1]. The maximum conductance *J*_*ij*_(*t*) determines the current transferred at the synaptic connection from the *j*th neuron to the *i*th neuron. The maximum conductance *J*_*ij*_(*t*) depends on the environmental random variables thus its dynamical model is represented as an independent diffusion process this mean value 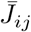 and standard deviation 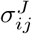 such that,

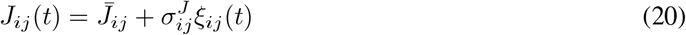

where *ξ*_*ij*_(*t*)*dt* = *dB*_*ij*_(*t*) are independent zero mean unit variance white noise processes derived from *B*_*ij*_(*t*) (independent standard Brownian motion).

We consider a plane network of *N* interacting neurons of the described dynamical and synaptic properties given by eqns (15) and (16) whose neuronal state of an individual neuron in the network (labeled as *αβ*) are identified by the vector (*u*_*αβ*_, *w*_*αβ*_, *y*_*αβ*_). For a network of partially connected neurons where the of connection for each neuronal unit is identified by a maximum spatial distance of influence Ω between neurons, the neuronal state is the solution of the following system of couple stochastic differential equations:

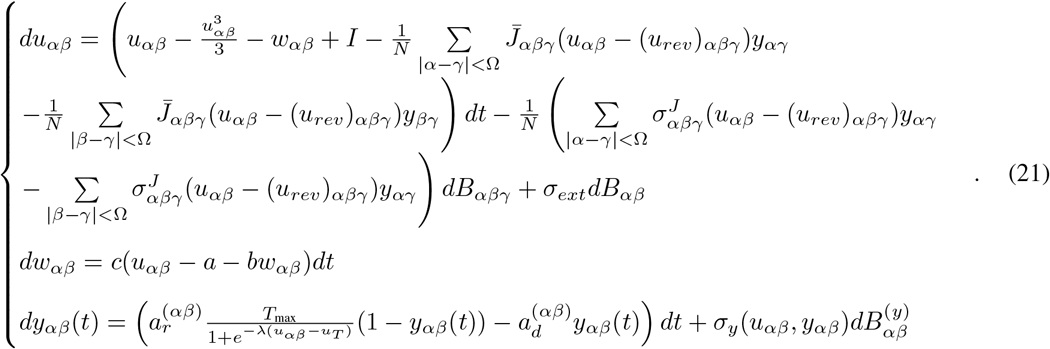

The solution to the system (21) constitutes the response of an array of neurons in the form of a two-dimensional grid of *N* × *N* neurons, each of which are connected to their neighboring neurons up to the *C*^th^ neuron apart from them in each direction. The system (21) constructs the small scale view of the neuronal network with nonlinear coupling between the nearby neurons.

### 3.2 Large-scale neural network dynamics

In the large-scale view of the model, we considers a network of neural populations by a system of reaction-diffusion cellular neural network (CNN) [Chan and Yang 1986]; a system of nonlinear dynamical cells coupled to their adjacent by a linear synaptic law approximating the spatial diffusion term (Laplacian). This takes the form of a system of spatio-temporal FHN equations. By analogy to the model discussed by Slavova and Zecca [2003], we consider a typical grid of a two-dimensional CNN (as shown in Fig. 1), to derive a two-dimensional formulation of the system.

**Figure 1.**
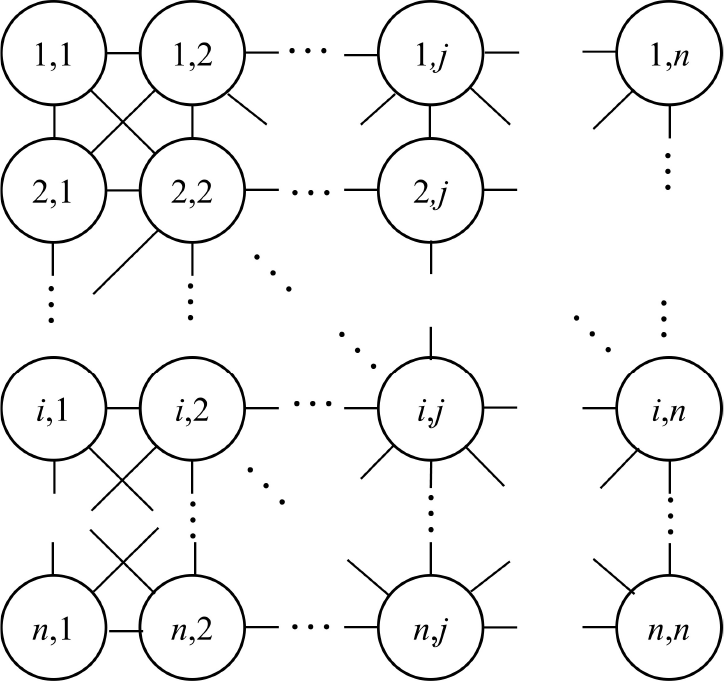
A typical cellular neural network design in 2D

**Figure 2.**
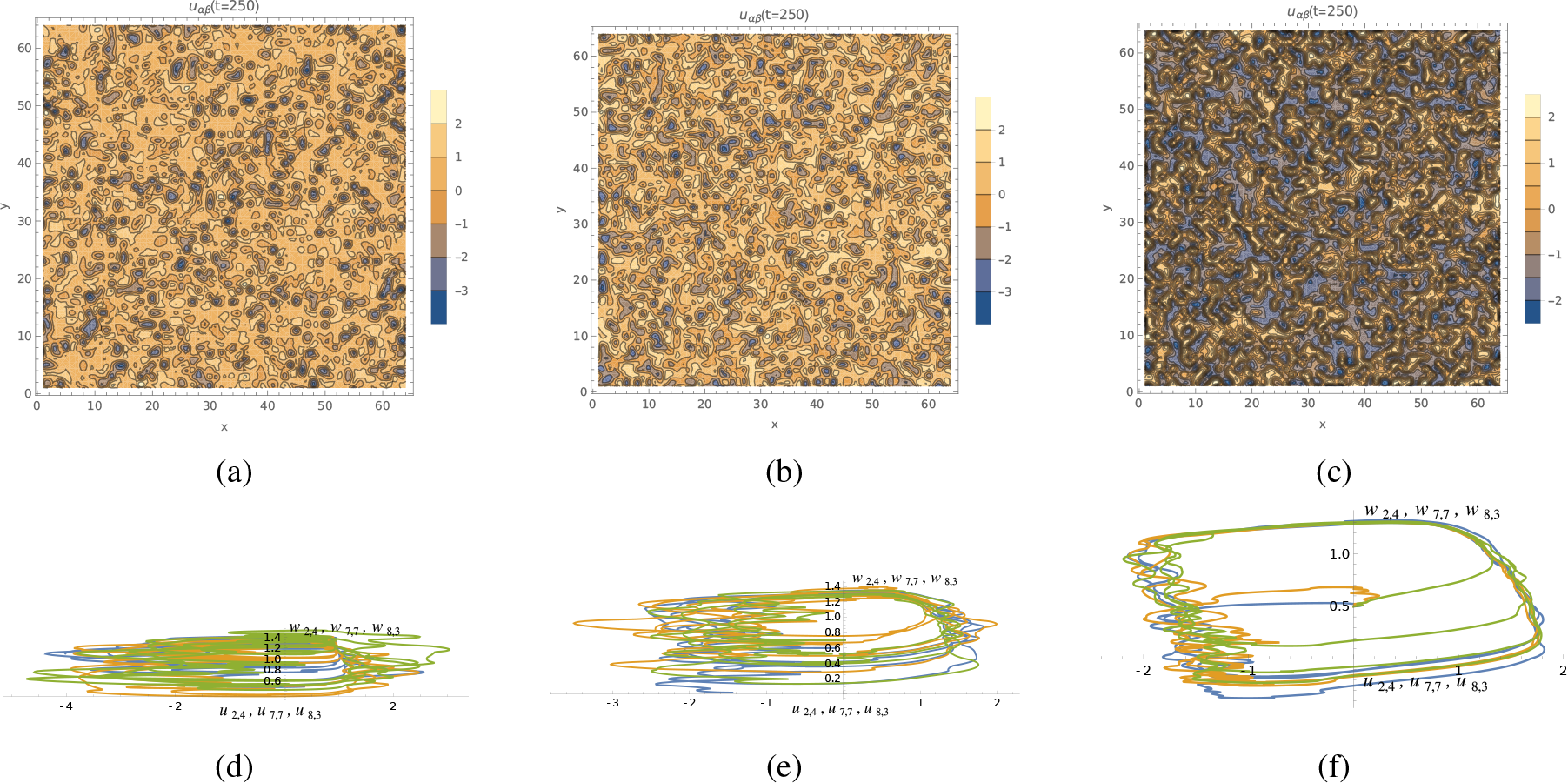
(a)-(c): voltage variations across the fine-scale view of the interconnected neural network. (d)-(f): orbits of voltage and slow variables at three chosen neurons (nodes) in the corresponding phase plane

**Figure 3.**
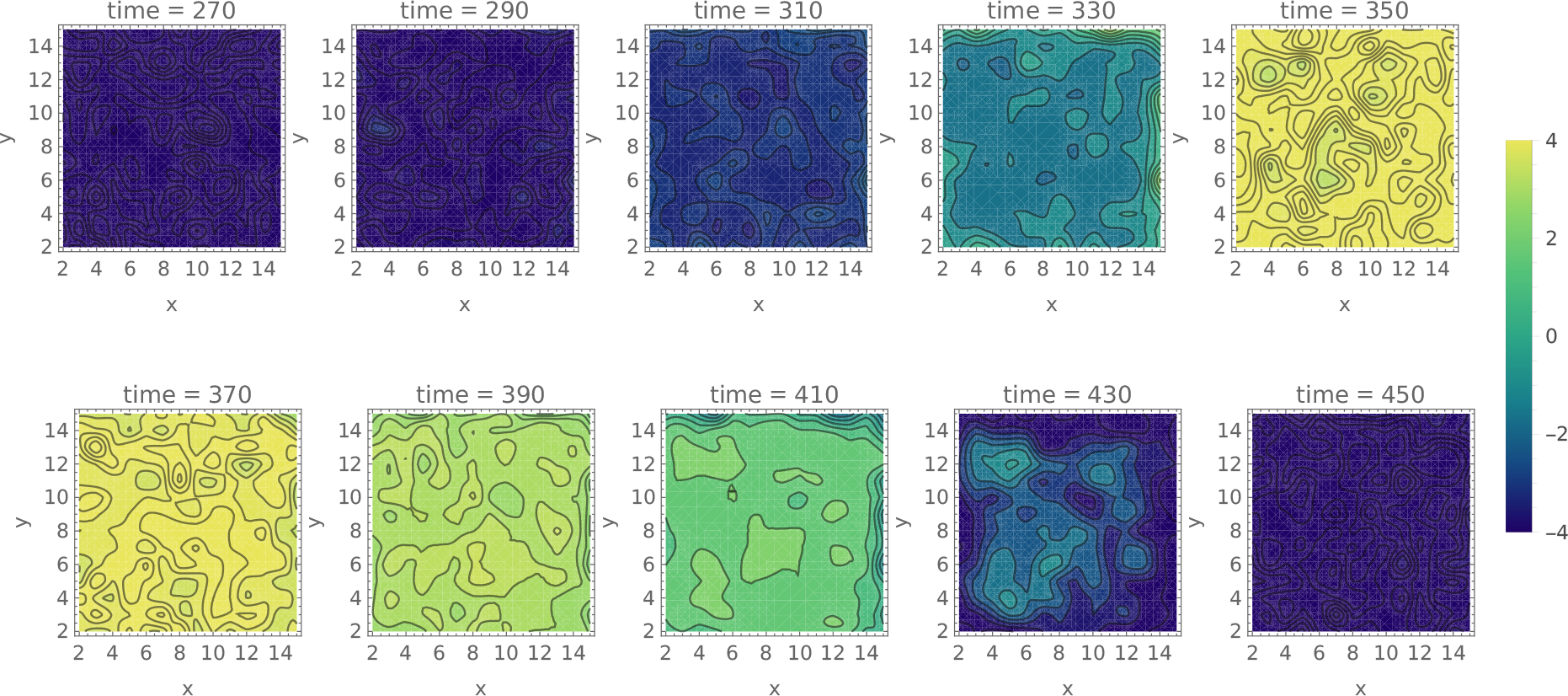
Wave evolution for the system of strong connectivity (fully connected network)

**Figure 4.**
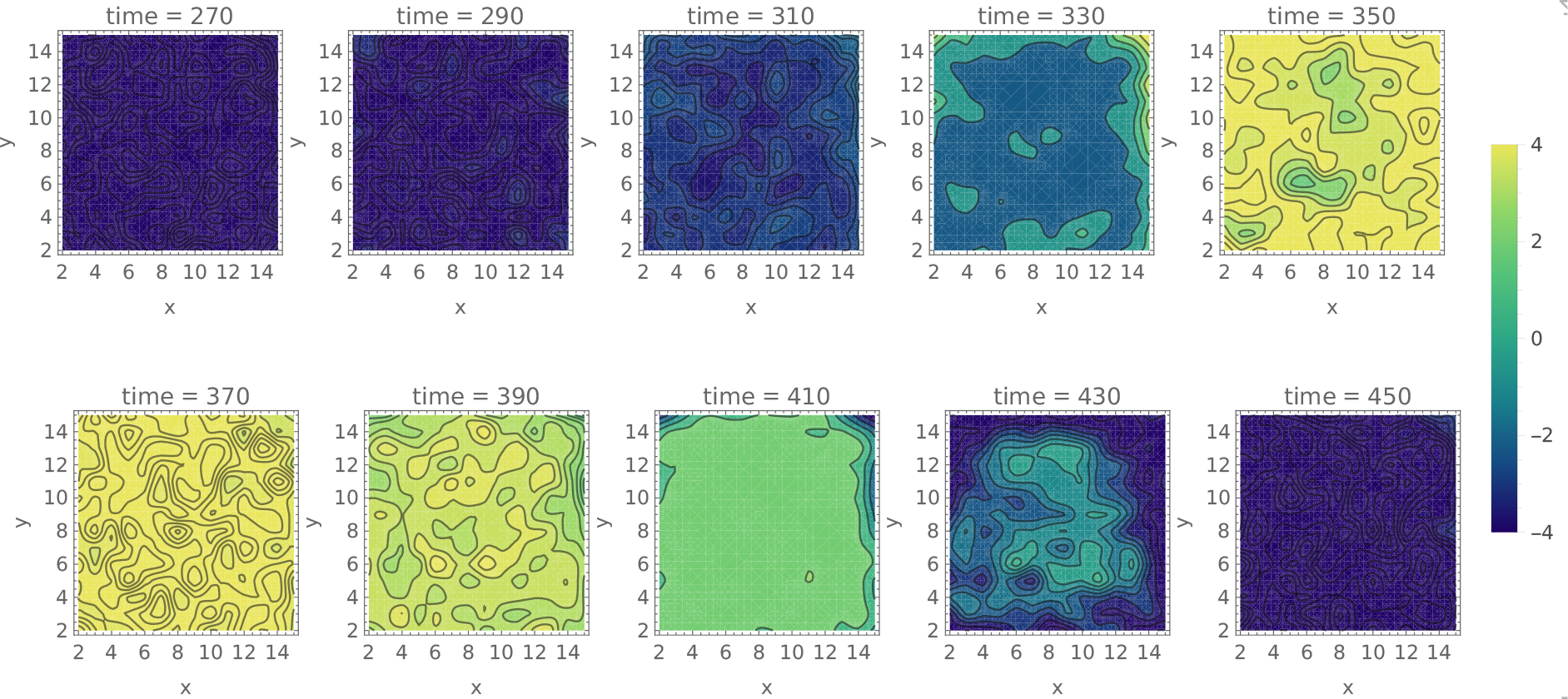
Wave evolution for the system of medium connectivity (each neuron connected to 20 closest neighbors)

**Figure 5.**
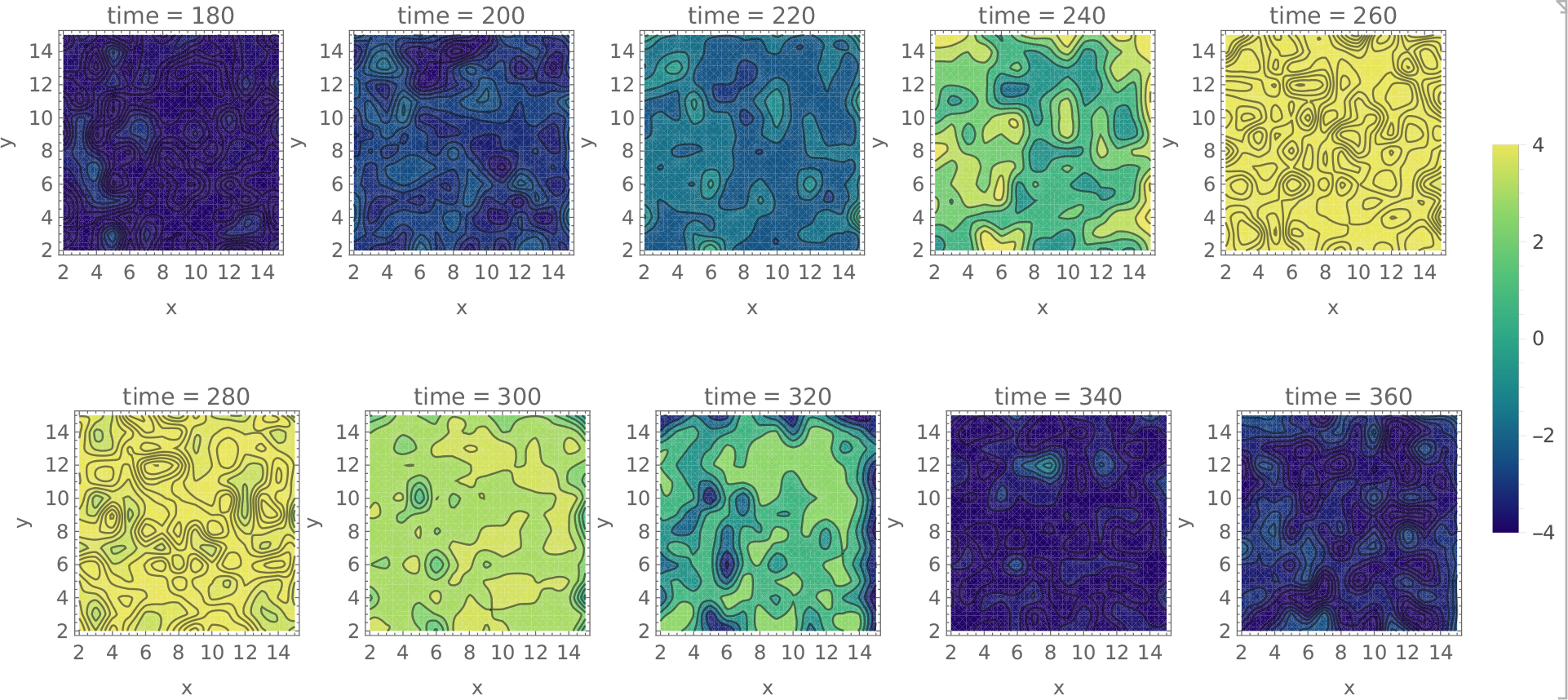
Wave evolution for the system of weak connectivity (each neuron connected to 3 closest neighbors)

Each circle in Figure 1 is a circuit unit representing a neuronal population and the links indicate coupling between the adjacent cells. The 2D FHN reaction-diffusion equation reads,

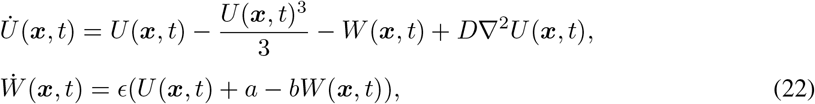

analogous to Eqn. (10), with the spatial derivative term amplified by a diffusion coefficient in two-dimensions where the Laplacian 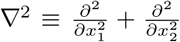, for a generic Cartesian point ***x*** = (*x*_1_, *x*_*2*_). The CNN model discussed in Slavova and Zecca [21] discretizes *u* and *w* into variables *U*_*ij*_(*t*), and *W*_*ij*_(*t*) assigned to each spatial grid point (*i, j*) in the network. Thus by the difference representation of the second partial derivative operators of the Laplacian ∇^2^*u* is approximated by,

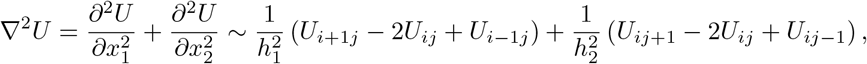

where *h*_1_ = Δ*x*_1_ and *h*_2_ = Δ*x*_2_, indicate the grid size in two directions, and *i* and *j* indicate the cell position in the CNN grid. Consequently, a FHN reaction-diffusion CNN system can be discretized as

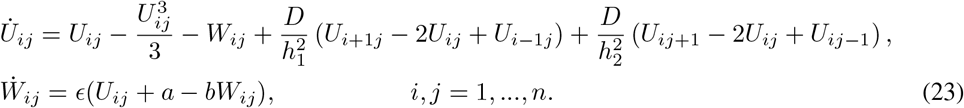

The system (23) is a system of nonlinear ordinary differential equations with linear coupling (synaptic law) characterizing the dynamical state of the CNN of *n* cells. For analysis of the current CNN system we take the periodic boundary conditions specifying that the values of *u* at the boundary points equal those of the opposite sides of a virtual extension of the grid from all sides. That is,

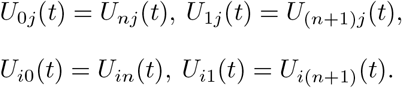

### 3.3 Between-scale coupling

The multi-scale architectural approach [2] that we adopt in this paper decomposes the neural system into a hierarchy of multi-scale evolution equations (known as between scale coupling) along with a within-sale coupling scheme to form an ensemble of nonlinear oscillator. Consider the solutions of the equations (23) and (21), i.e., the vectors ***U*** = {*U*_*ij*_, *W*_*ij*_}, and ***u*** = {*u*_*αβ*_, *w*_*αβ*_} denote two ensembles of a network variables at two spatial scales *j*_1_ and *j*_2_, corresponding respectively, to the course and the fine scales. In reference to the discussion leading to eqn (26), 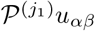 and 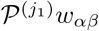 are defined as the projection of the fine-scale solutions *u*_*αβ*_ and *w*_*αβ*_ from the scale *j*_2_ onto the course scale *j*_1_. In the forthcoming numerical evaluations we assign scale indices *j*_1_ = 4 and *j*_2_ = 0. The dynamics of the course scale system with the fine-scale spatial coupling effects is now described by ([2])

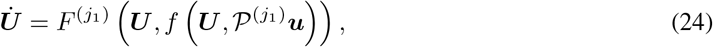

where function *f* specifies the inter-scale coupling rule. Breakspear Stam [2] consider a linear inter-scale coupling by introducing a linear combination of the course scale dynamics by a factor *G* of the difference between the course scale response and the fine-scale projection onto it. Effectively, the projection 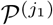 filters the spatial variance of the fine-scale results to inject the relevant information from the finer resolution *u*_*αβ*_ and *w*_*αβ*_, to the course scale responses *U*_*ij*_, *W*_*ij*_. In addition to the disruptive effect of the projection projection 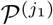, the mean field of the fine scale response is also linearly combined with the course scale dynamics to account for the global synchronized effects of the fine-scale response. The final scale-specific adjustment corresponds to the difference in the temporal dynamics at each scale. This is introduced in the system by time-scale multipliers.

Putting everything together, the multi-scale dynamics of the CNN incorporating the effects of the finescale stochastic dynamics of interacting neurons is written as follows:

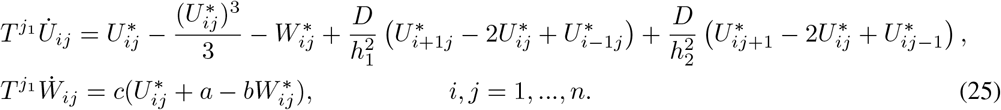

where

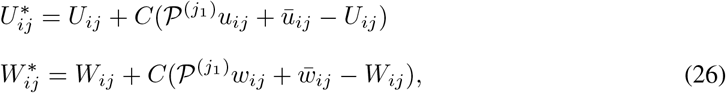

and *T* is a time scale multiplier specifying the internal time scale corresponding to each spatial scale level. The functions *u*_*ij*_ and *w*_*ij*_ correspond to the scale equivalent nodes in the fine-scale view. For instance, if two successive scales differ by a factor of four 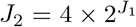 then equivalently each *u*_*αβ*_ equals *u*_4*i*4*j*_. The bar over 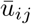 and 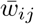 indicates the spatial average of the functions over the network domain at any given time. See Breakspear and Stam [2] for a comprehensive account of the multi-scale scheme on which the current model is founded. In the coming section we provide a numerical example that demonstrates the solution form of the multi-scale neural network.

## 4 Numerical Example

We present a numerical simulation of the equation (25) in this section. Consider a course scale view with *J*_2_ = 4 and a fine scale view with *J*_1_ = 0. For the numerical analysis, we employ the Shannon scaling and wavelet basis functions as they provide an adequate frequency localization, although the choice of the wavelet bases is arbitrary. We construct a two-tiered structure of the course scale 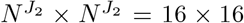 view of the plane, and the fine scale view 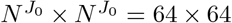. Three fine scale within-scale connectivity scenarios are investigated. The fist scenario corresponds to a weak within scale coupling wherein the neural units at the fine scale are effectively connected to the 3 nearest neighbouring neurons (Ω = 3 in eq. (21)). The second scenario assumes a medium strength connectivity where Ω = 20. Finally we consider a fully connected fine scale network where each neural unit is connected to all the other neurons in the plane. For the simulation purpose, we adopt the numerical values from Baladron et al [1] presented in table (1)

**Table 1:**
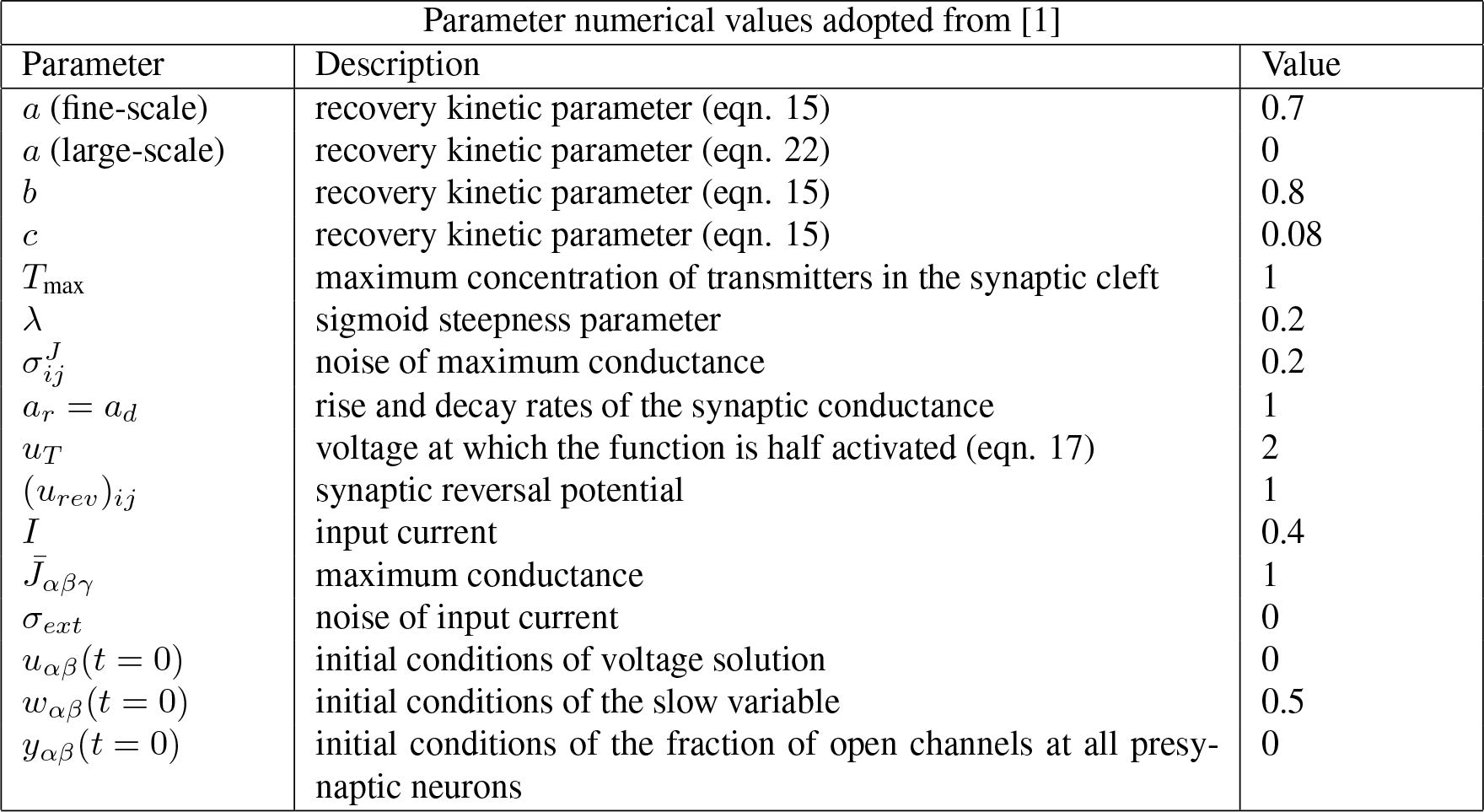
Numerical values for the involved parameters.

A sample result of the fine scale solutions for the three connectivity scenarios are presented in figure (2), where we illustrate the spatial variations of the normalized voltage *u*_*αβ*_ illustrated in a snapshot at time *t* = 250, as an example. Clearly, for the short connectivity of neural ensemble, there are higher spatial variations with highly localized nature of the response all over the plane, whereas, for a fully connected network the solution characterizes with a more uniform distribution and sparsely random localizations. Sample limit cycles of the stochastic solutions corresponding to the three scenarios are illustrated in the same figure (2), and quite obviously the stronger neural connectivity intensifies the associated noise in the neuronal system. For the multi scale simulation, we set the coupling coefficient to *C* = 0.5. The connectivity effects of the underlying fine scale neural mass dynamics on the large scale network of reaction diffusion kind is illustrated in Figures (3-5). As observed, for the fully (or moderately) connected fine scale network the relatively synchronized global background in the noisier medium (as a result of the stronger connectivity) give rise to high definition travelling front. This clear cut wave front is more uniform for the moderate connectivity where each neuron is connected to 20 closest neighbours, whereas the higher noise for the fully connected network downplays the synchronous effect on the global emergence of the travelling waves. This is due to the fact that the mean field synchronous voltage variations and the involved noise cancel each other’s effect to amount to a nearly vanishing spatial average at any time frame. On the other hand, for the weakly connected fine scale neural network where each neuron is connected with only 3 closest neighbours, the highly localized and intense variations in the presence of less network voltage noise generate multiple disordered travelling wave fronts in the large scale system. It must be mentioned that in all cases, we set the large scale reaction diffusion coefficient in two directions 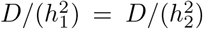 to 0.05 so that at the scale and with the chosen network parameters under scrutiny the travelling waves appear as an observable phenomenon. There is indeed ranges of reaction-diffusion coefficients in which an overall asynchronous or synchronous large scale behaviour can appear as a dominant pattern. Moreover, it should be noted that we limit our numerical evaluation of the FHN multi scale construct to the voltage variable and, thus we ignore the interdependence of the slow variable between different scales in the current study.

## 5 dynamic behaviour of the multi scale architecture

While in the previous section we evaluated the model and observed the influences through the numerical measurement, a closer investigation of the model itself can provide a better understanding of the behaviour of the system. In this section, we perform an approximate evaluation of the dynamical system of the reaction-diffusion model under the projection of the fine-scale functional connectivity of the underlying neural units. The multi-scale model described by eq. (25) can be characterized as a spatially discrete stochastic system where the parameters of the dynamical system and the input are stochastic processes determined by the fine-scale output. The properties of the processes involved are determined by the types of Fokker-Planck equations that result from the original fine-scale stochastic differential equations [1, 18]. The deterministic part of the equation (25), however, behave dynamically similar to the CNN model in [21]. In this paper, we set our focus to the simplified deterministic part of the large-scale equation which incorporates the between-scale coupling coefficient as well as the scale specific time factor. A more comprehensive analysis of the stochastic large-scale equation fully incorporating the fine-scale effects will be the topic of a future study. First, we would like to establish the harmonic balance argument for the deterministic part of the equation (25), i.e.

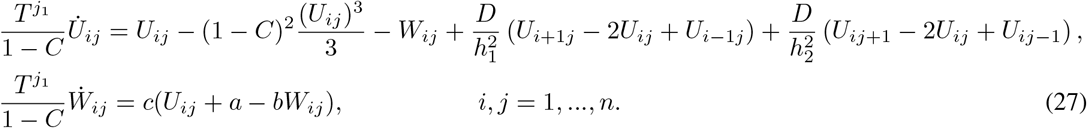

By the harmonic balance principle we can transform the equation to a quasi-linear representation [22]

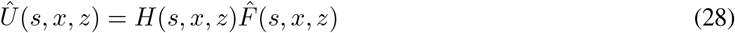

with the linear part of the system to construct the transfer function

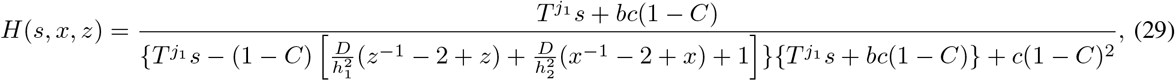

that relates *Û*and 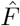, respectively indicating the transformed form of *U* and the nonlinear part 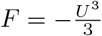 by the following continuous-time discrete space Fourier transform:

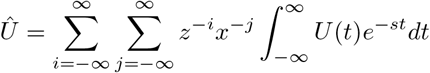

where, *z* = exp(ı*γ*_*z*_), *x* = exp(ı*γ*_*x*_), 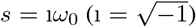 and the solution is approximated by the harmonic function in the form of a periodic wave as

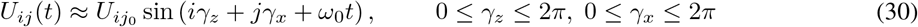

Once we apply the periodicity of the boundary conditions to (30), we obtain 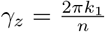 and 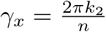, with natural numbers 1 ≤ *k*_1,2_ ≤ *n*. Further, we approximate the systems output by its fundamental component of the Fourier expansion,

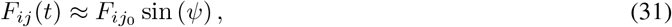

where *ψ* = *iγ*_*z*_ + *jγ*_*x*_ + *ω*_0_*t* designates the argument of *sine* and 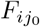 can be determined through the Fourier expansion as

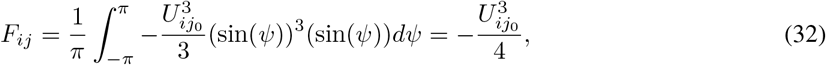

We incorporate the values *z* = exp(ı*γ*_*z*_), *x* = exp(ı*γ*_*x*_), *s* = ı*ω*_0_ into the expression for the transfer function (29) to have,

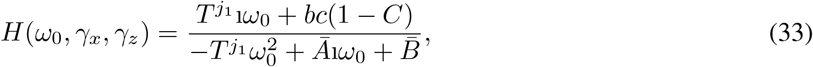

where

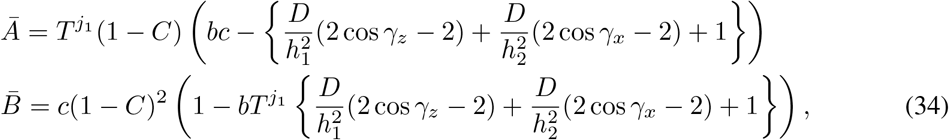

The harmonic ansatz of the approximated transformed nonlinearity (31) dictates the constraints,

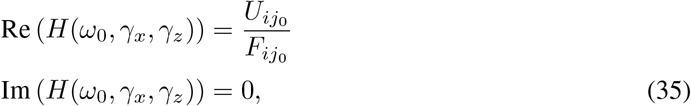

whence we can solve (35) for 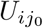 and *ω*_0_. Thus, we find an approximation for the periodic wave solution with the amplitude,

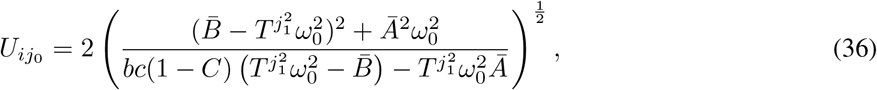

and the frequency,

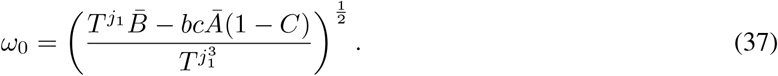

Therefore, similar results to those mentioned by Slavova and Zecca [21] is acquired which imply the approximate conditions under which such periodic solutions can be expected as well as the level of accuracy of the approximation.

Next, we are interested in evaluating the deterministic part of the multi-scale equation (27) in terms of the conditions under which travelling wave solutions exist. The existence and stability of the FHN model in its spatio-temporal form (PDE) have been subjects of extensive studies. To name a few, Jones [14] use Evans approach to prove the stability of the FHN travelling waves, Deng [6] demonstrated the existence of multiple travelling fronts and backs for the FHN equation of bistable type, Yanagida [25] used an eigenvalue analysis to prove that the FHN equation travelling fronts are asymptotically stable under some given requirements. Relatively more recently attentions have been attracted to the problems of existence and stability of waves in systems of coupled nonlinear ordinary differential equations. The earlier studies [5, 10, 23] determined the conditions of the existence of wave fronts in discrete coupled Nagumo systems. More recently Hupkes, Sandstede and Schouten-Straatman [12, 13, 19, 20] investigated the existence and stability of FHN travelling waves in a discrete lattice system whose diffusion term appears in a difference form and discussed the caveats which concern the emergence of wave fronts and pulses. Here in our brief assessment, we are concerned with the possibility of appearance of travelling wave solutions in the following form,

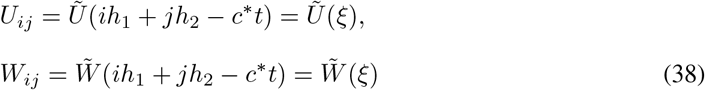

where *c*^*^ *>* 0 is the velocity of the wave. We substitute (39) into (27) and to further simplify the argument, consider a uniform lattice with *h* = *h*_1_ = *h*_2_. We obtain the following differential-difference equation,

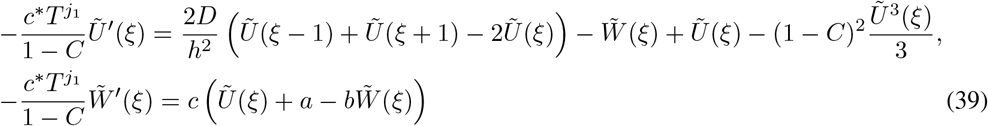

The fixed points of the system (39) (the solutions of the stationary problem) satisfy the equation (39) with 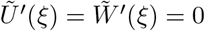 0 and 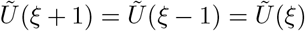. Under such circumstances as well as the following condition,

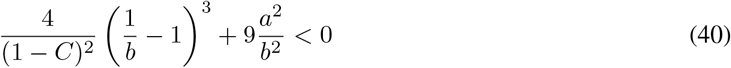

we find the three fixed point in the phase plane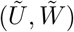, namely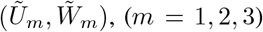, where by the Vieta’s formula,

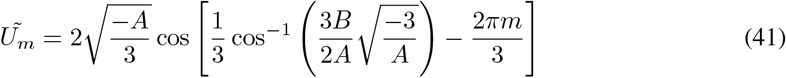

and

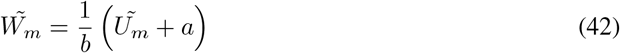

To investigate the stability conditions of the stationary solutions we linearize the system (39) about the determined fixed points. The stability of the fixed points are determined by solving the following eigenvalue problem:

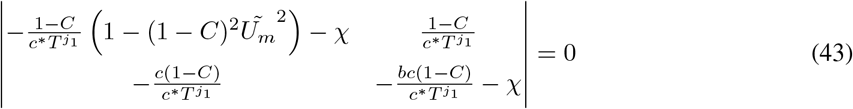

which can be translated into the characteristic equation,

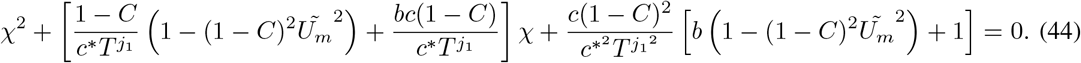

The fixed points 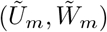 for which 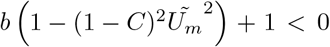 are saddle point meaning that the system trajectories approach them on a stable manifold and depart from them on an unstable manifold. The points 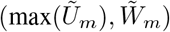 and 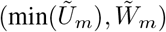 with the maximum and the minimum *Ũ*_*m*_ require to have the saddle point conditions and the trajectory must remain in *Ũ* ∈ min(*Ũ*_*m*_), max(*Ũ*_*m*_) to create a heteroclinic orbit. Now having the following boundary conditions,

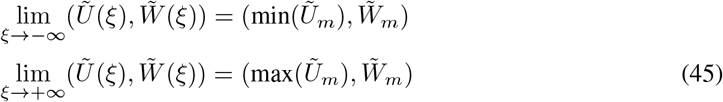

or

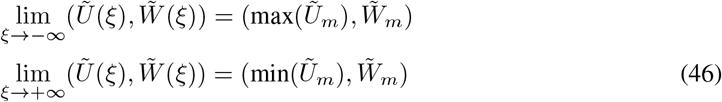

we can show that the solution 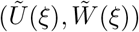 of the deterministic part of the multi-scale system corresponds to a travelling wave front (or back). Further more thorough analysis of the complete multi-scale stochastic system is required to gain a more accurate perspective of the behaviour of the model. It i

## 6 Conclusions

In this paper, we adopted the multi-scale architecture based on the wavelet decomposition scheme introduced by Breakspear and Stam to design a two-tiered model incorporating the stochastic interaction of individual neurons to the large-scale view of the reaction diffusion system. Through this study we evaluated the implications that arise from such multi-scale construct in terms of the emergence of travelling waves under the certain connectivity conditions of the underlying fine-scale network as well as the diffusion coefficient measured at the large scale. It is obvious, however, that he connectivity of individual neurons affect the largescale diffusion coefficient which is itself a measure of intra-scale coupling between the neural units. The numerical analysis demonstrates that the neural synchronization mode which is most suited to the emergence of travelling waves occurs as a result of medium level of connectivity strength of the underlying fine-scale network. We ultimately presented an approximate evaluation of the resulting dynamical system to determine the conditions under which travelling wave phenomena may be expected. Obviously, our simplified analysis demonstrates that the multi-scale construct which depends on between-scale coupling and scale-specific time parameter alters the conditions of the travelling wave solution and a higher coupling coefficient leads to more limited conditions for the appearance of travelling wave fronts in the large-scale system. However, the problem of determining those conditions remains open to further studies through the available techniques of spectral function analysis, Fredholm alternative machinery and the exchange lemma, etc. The numerical simulation measurements juxtaposed with the results from the large-scale reaction diffusion system evidence the possibility of travelling waves in accordance with the connectivity strengths (which determine the stochastic stimulus of the large-scale system), the spatial diffusion and other parameters of the system which determine the state trajectories.

## Acknowledgements

The authors thank the Canadian Institutes of Health Research for the support through a post-doctoral fellowship award.

## References

[1] J. Baladron, D. Fasoli, O. Faugeras, and J. Touboul. Mean-field description and propagation of chaos in networks of hodgkin-huxley and FitzHugh-nagumo neurons. The Journal of Mathematical Neuroscience, 2(1), may 2012.

[2] M. Breakspear and C. J. Stam. Dynamics of a neural system with a multiscale architecture. Philosophical Transactions of the Royal Society B: Biological Sciences, 360(1457):1051–1074, may 2005.

[3] Michael Breakspear. Dynamic models of large-scale brain activity. Nature Neuroscience, 20(3):340–352, feb 2017.

[4] Richard B. Buxton. Introduction to functional magnetic resonance imaging. Cambridge University Press, 2009.

[5] Shui-Nee Chow, John Mallet-Paret, and Wenxian Shen. Traveling waves in lattice dynamical systems. Journal of Differential Equations, 149(2):248–291, nov 1998.

[6] Bo Deng. The existence of infinitely many traveling front and back waves in the FitzHugh–nagumo equations. SIAM Journal on Mathematical Analysis, 22(6):1631–1650, sep 1991.

[7] FitzHugh. Mathematical models of threshold phenomena in the nerve membrane. Bull. Math. Biophysics, 17:257–278, 1955.

[8] R. Fitzhugh. Theoretical effect of temperature on threshold in the Hodgkin-Huxley nerve model. Journal of General Physiology, 49(5):989–1005, may 1966.

[9] R. FitzHugh. Mathematical models of excitation and propagation in nerve. In Biological Engineering, chapter 1, pages 1–85. McGraw-Hill Book Co., 1969.

[10] D. Hankerson and B. Zinner. Wavefronts for a cooperative tridiagonal system of differential equations. Journal of Dynamics and Differential Equations, 5(2):359–373, apr 1993.

[11] A. L. Hodgkin and A. F. Huxley. A quantitative description of membrane current and its application to conduction and excitation in nerve. The Journal of Physiology, 117(4):500–544, aug 1952.

[12] H. Hupkes and B. Sandstede. Stability of pulse solutions for the discrete FitzHugh–nagumo system. Transactions of the American Mathematical Society, 365(1):251–301, jul 2012.

[13] H. J. Hupkes and E. S. Van Vleck. Travelling waves for complete discretizations of reaction diffusion systems. Journal of Dynamics and Differential Equations, 28(3-4):955–1006, jan 2015.

[14] Christopher K. R. T. Jones. Stability of the travelling wave solution of the FitzHugh-nagumo system. Transactions of the American Mathematical Society, 286(2):431–469, 1984.

[15] C. Morris and H. Lecar. Voltage oscillations in the barnacle giant muscle fiber. Biophysical Journal, 35(1):193–213, jul 1981.

[16] J. Nagumo, S. Arimoto, and S. Yoshizawa. An active pulse transmission line simulating nerve axon. Proceedings of the IRE, 50(10):2061–2070, oct 1962.

[17] Paul L. Nunez and Ramesh Srinivasan. Electric Fields of the Brain. Oxford University Press, jan 2006.

[18] Hannes Risken. The Fokker-Planck Equation. Springer, Berlin Heidelberg, 1996.

[19] Willem M. Schouten-Straatman and Hermen Jan Hupkes. Traveling waves for spatially discrete systems of FitzHugh–nagumo type with periodic coefficients. SIAM Journal on Mathematical Analysis, 51(4):3492–3532, jan 2019.

[20] W.M. Schouten-Straatman and H.J. Hupkes. Travelling wave solutions for fully discrete FitzHugh-nagumo type equations with infinite-range interactions. Journal of Mathematical Analysis and Applications, 502(2):125272, oct 2021.

[21] A. Slavova and P. Zecca. CNN model for studying dynamics and travelling wave solutions of FitzHugh–Nagumo equation. Journal of Computational and Applied Mathematics, 151(1):13–24, feb 2003.

[22] M. Vidyasagar. Nonlinear Systems Analysis (Classics in Applied Mathematics). SIAM: Society for Industrial and Applied Mathematics, 2002.

[23] Erik S. Van Vleck, John Mallet-Paret, and John W. Cahn. Traveling wave solutions for systems of ODEs on a two-dimensional spatial lattice. SIAM Journal on Applied Mathematics, 59(2):455–493, jan 1998.

[24] E. Weinan. Principles of multiscale modeling. Cambridge University Press, 2011.

[25] E. Yanagida. Stability of travelling front solutions of the fitzhugh-nagumo equations. Mathematical and Computer Modelling, 12(3):289–301, 1989.

